# Integrated *in silico* identification of fungal-derived dual-target inhibitors for anti-schistosomal drug discovery

**DOI:** 10.64898/2026.09.04.749322

**Authors:** Tamal Paul, Md. Al-Amin, Uma Shill, Nowshin Tabassum, Kaniz Fatema, Easin Al Riad, Milon Kumar Sarkar, Ahmad Abdullah Mahdeen, Ananya Majumder, Aminun Naher, Fubliha Firose Auishe, Fariha Bushra Khan, Foisal Ahmad, Hossain Bin Afridy, Md. Ashik Iqbal, Arnob Biswas, Khandokar Fahmida Sultana, Swagato Dutta, Sabrin Bashar

**Affiliations:** Department of Microbiology, Noakhali Science and Technology University, Noakhali-3814, Bangladesh; Department of Chemistry, Jagannath University, Dhaka-1100, Bangladesh; Department of Genetics and Plant Breeding, Bangabandhu Sheikh Mujibur Rahman Agricultural University, Gazipur-1706, Bangladesh; Department of Pharmacy, Noakhali Science and Technology University, Noakhali-3814, Bangladesh; Department of Microbiology, Notre Dame University Bangladesh (NDUB), Dhaka-1000, Bangladesh; Department of Microbiology, Jahangirnagar University, Dhaka-1342, Bangladesh; Department of Biochemistry and Molecular Biology, Jahangirnagar University, Dhaka-1342, Bangladesh; Department of Microbiology, Primeasia University, Dhaka-1213, Bangladesh; Department of Pediatrics, University of Alberta, Edmonton, AB T6G 1C9, Canada

**Keywords:** *Schistosoma mansoni*, fungal metabolites, molecular docking, MD simulation, DFT, QSAR, drug discovery

## Abstract

Praziquantel, which has limited efficacy against young parasites and reduces susceptibility, is the principal therapy for schistosomiasis, a neglected tropical illness. This work identified fungal-derived compounds with potential dual inhibitory action against *Schistosoma mansoni* DHODH and cathepsin B1 using an integrated computational method. From 1,831 fungal metabolites, 120 were selected for molecular docking after structure standardisation, drug-likeness evaluation, ADMET, and toxicity screening. Ganoderlactone B (CID_122184973) and Spiroapplanatumine F (CID_132962217) strongly bound to DHODH and CB1, with docking affinities of −10.6 and −10.0 kcal/mol and -9.0 and -8.9, respectively. Docking validation showed good discrimination between active compounds and decoys with area-under-the-curve values of 0.98 for DHODH and 0.96 for CB1. The protein-ligand interaction study showed hydrogen bonding, hydrophobic interactions, and van der Waals contacts at the binding sites. In 100 ns molecular dynamics simulations, protein complexes exhibited modest backbone deviations and maintained compactness, although CID_132962217 showed sustained binding and a constrained conformation. MM-PBSA analysis identified CID_122184973 as the best common ligand for both targets. At the B3LYP/6-31G level, density functional theory calculations showed that CID_139584993 had the lowest HOMO-LUMO energy gap and hardness values of 3.552 and 1.776 eV, respectively, and the highest softness value of 0.282 eV^-1^, indicating greater electronic reactivity. Electrostatic potential mapping and QSAR predictions supported their interaction and antiparasitic potential. Overall, CID_122184973 and CID_132962217 are promising fungal scaffolds for dual-target anti-schistosomal drug development. Experimental enzyme inhibition, parasite viability, toxicity, and *in vivo* research are needed to confirm computational findings.

## 1. Introduction

Schistosomiasis is one of the most important parasitic infections affecting human health worldwide and is currently reported to affect ∼218 million people in tropical and subtropical regions of Africa, the Middle East, South America, and Southeast Asia [1]. This neglected tropical disease (NTD) is associated with significant morbidity and mortality, with over 90% of the global burden occurring in sub-Saharan Africa [2,3]. The clinical course of schistosomiasis includes an acute and a chronic phase. Acute schistosomiasis, also known as Katayama fever, usually occurs within 4-6 weeks of infection and is characterized by fever, malaise, and eosinophilia. Chronic disease develops subsequently and may manifest as dry cough, mild fever, weight loss, and anaemia [4]. Long-term infection with schistosomiasis causes significant morbidity, such as hepatic fibrosis, portal hypertension, bladder pathology, kidney dysfunction, growth retardation and enhanced risk of bladder cancer and hence a significant socioeconomic burden on endemic communities [1,5].

The three species responsible for human schistosomiasis (*Schistosoma mansoni*, *Schistosoma haematobium*, and *Schistosoma japonicum*) are distributed globally, with *Schistosoma mansoni (S. mansoni)* being the most common responsible agent in Africa, Yemen, and Brazil [1,6]. *S. mansoni* has a complex life cycle that alternates between humans, the definitive host, and freshwater snails of the genus *Biomphalaria*, which serve as the intermediate host. Transmission is by contact with contaminated water containing the infective stage, the cercariae, which penetrate the skin and mature into adults in the mesenteric vasculature [4,6]. The pathological effects of *S. mansoni* infection are primarily associated with egg deposition in host tissues, which triggers a strong inflammatory response. The parasitic eggs, particularly in the liver and intestine, elicit T helper 2 (Th2) cell-mediated granulomatous reactions leading to portal fibrosis, presinusoidal portal hypertension, and hepatosplenomegaly [7]. This renders *S. mansoni*-induced schistosomiasis haematobium a major cause of morbidity in endemic regions, and the need for effective therapeutic strategies is therefore of the utmost urgency.

The survival and proliferation of *S. mansoni* depend critically upon efficient pyrimidine metabolism and protein processing. Dihydroorotate dehydrogenase (DHODH) is a mitochondrial enzyme catalyzing the fourth step of de novo pyrimidine synthesis, converting dihydroorotate to orotate, a rate-limiting step essential for nucleotide biosynthesis and parasite replication [8,9]. *Schistosoma mansoni*’s capacity to salvage preformed pyrimidines is limited and poorly characterized, leaving the parasite primarily reliant on de novo pyrimidine synthesis to meet its nucleotide demands, thereby making DHODH a promising therapeutic target. Inhibition of DHODH leads to depletion of pyrimidine pools, impairing DNA replication and triggering parasite death [10–12]. Complementary to this metabolic vulnerability, cathepsin B1 (CB1) is a major cysteine protease abundantly expressed in *S. mansoni* and plays a critical role in host immune evasion, nutrient acquisition, and hemoglobin degradation [10,13]. CB1 facilitates the degradation of host immunoglobulins and contributes to the parasite’s ability to evade antibody-mediated immunity, rendering it indispensable for parasite survival within the hostile host environment [14–16]. Furthermore, CB1 participates in the processing of parasite antigens, influencing the host’s Th response and modulating granulomatous inflammation [17].

The therapeutic options for schistosomiasis remain limited, with praziquantel (PZQ) serving as the cornerstone of treatment and the only World Health Organization (WHO)-recommended drug for more than three decades [18]. PZQ is very effective against adult worms, but there are some important limitations. It is not as effective against immature schistosomes as it is against adults; it may need to be repeated if transmission is high, and there have been reports of reduced susceptibility and the emergence of resistance in certain endemic areas, such as Egypt and Senegal [19,20]. Other alternatives, like oxamniquine, have limited activity and are limited in use due to safety concerns and geographic specificity, making them less useful, especially in resource-limited environments. The absence of effective preventive interventions, continued transmission in endemic areas and regular reinfections underscore the need for new anti-schistosomal drugs that are more effective, less toxic, active against multiple parasite developmental stages and have novel mechanisms of action to combat emerging drug resistance and facilitate sustainable disease control [20,21].

Natural products remain one of the most fertile sources of therapeutic agents. Fungal secondary metabolites are a source of bioactive molecules with antifungal, antiparasitic, anticancer, antiviral, and anti-inflammatory properties [22]. Fungi can produce structurally complex molecules such as polyketides, terpenoids, alkaloids, peptides, and hybrid metabolites, many of which have novel pharmacological properties. Many fungal metabolites have been shown to exhibit highly attractive antiparasitic activity against protozoan and helminth parasites and have thus been proposed as potential scaffolds for antiparasitic drug development [23]. However, there is a lack of studies on the potential of fungal metabolites to target molecular pathways associated with schistosomiasis.

In recent years, computer-aided drug discovery (CADD) has continually improved the discovery of potential drug candidates with less time and expense, becoming a powerful tool. Computational methods like molecular docking, molecular dynamics (MD) simulation, density functional theory (DFT), quantitative structure-activity relationship (QSAR) analysis, and *in silico* ADMET profiling can provide a comprehensive evaluation of the binding of a ligand to the target receptor, molecular stability, electronic properties, biological activity, and pharmacokinetic behavior before any experimental validation [24,25]. These strategies combined offer a powerful tool for prioritizing compounds of therapeutic interest.

This study employed an integrated computational pipeline to systematically screen a curated library of fungal-derived natural products for dual inhibitory activity against *S. mansoni* DHODH and CB1. Following the assessment of ADMET and drug-likeness properties, selected compounds were evaluated by molecular docking to identify high-affinity dual-target inhibitors.

The most promising protein-ligand complexes were further characterized through molecular interaction analysis, molecular dynamics simulations, density functional theory (DFT) calculations, molecular electrostatic potential mapping, and quantitative structure-activity relationship (QSAR)-based biological activity prediction. By integrating these complementary computational approaches, this study identifies promising fungal metabolites with potential anti-schistosomal activity and provides a rational framework for the future development of dual-target therapeutics against schistosomiasis.

## 2. Methods and materials

### 2.1. Protein selection and preparation

The protein structures 6UY4 (DHODH) and 4I07 (CB1) were selected as therapeutic targets due to their essential roles in *Schistosoma mansoni* survival and the availability of high-resolution crystal structures. DHODH is indispensable for de novo pyrimidine biosynthesis and possesses a *Schistosoma*-specific structural domain absent from the human homolog, thereby facilitating selective inhibitor design [10]. CB1 is the major gut-associated cysteine protease involved in host hemoglobin degradation, and its inhibition significantly impairs parasite viability, supporting its potential as an anti-schistosomal drug target [13].

The X-ray crystal structures of both proteins were retrieved from the RCSB Protein Data Bank (https://www.rcsb.org/). The 6UY4 and 4I07 form the same A chain with a sequence length of 379 AA (amino acids) and 254 AA, respectively. 6UY4 has a resolution of 2.80 Å and an R-free value of 0.246 [26], while 4I07 has a resolution of 1.30 Å and an R-free value of 0.177 [27]. The crystallographic water molecules were removed prior to docking to prevent the non-conserved solvent effect. The structures were then screened for missing residues, and the missing residues and gaps in the deposited model were reconstructed using AlphaFold Server (https://alphafoldserver.com/). All resulting models were visually examined to ensure that the backbone is connected and that the stereochemistry is accurate in the reconstructed regions before being used as a receptor for further computational analysis.

### 2.2. Ligand selection

Following target protein selection, a ligand library was compiled from the fungal metabolites database of medicinal fungi (https://cb.imsc.res.in/mefsat/). The selected compounds were then searched in the PubChem database (https://pubchem.ncbi.nlm.nih.gov/) to find a unique chemical identifier for each compound, and the PubChem compound identifier (CID) and SMILES were also recorded to ensure unambiguous annotation. Finally, the 3D structural records for each ligand were retrieved from PubChem to facilitate subsequent computational analyses.

### 2.3. ADMET prediction and drug-likeness screening

Antifungal metabolites with favourable pharmacokinetic and safety profiles were prioritised through *in silico* ADMET evaluation prior to docking and molecular dynamics analysis. SwissADME (https://www.swissadme.ch/) was employed to evaluate drug-likeness and key physicochemical descriptors, such as molecular weight (MW), XlogP, number of rotatable bonds, hydrogen bond donors (HBD), hydrogen bond acceptors (HBA), and overall compliance with Lipinski’s Rule of Five [18,28].

Computational toxicity profiling was conducted using pkCSM (https://biosig.lab.uq.edu.au/pkcsm/) to augment physicochemical screening with toxicity- and ADME-related risk predictions. The anticipated endpoints were cutaneous sensitisation, hepatotoxicity, and AMES mutagenicity. Furthermore, parameters related to absorption and distribution were recorded, with a particular emphasis on gastrointestinal (GI) absorption. Compounds predicted to have high blood-brain barrier (BBB) permeability were prioritised, while candidates predicted to be non-permeant were preferred to minimise the risk of Central Nervous System (CNS) exposure [29,30].

### 2.5. Molecular docking

A molecular docking evaluation was conducted to assess the binding affinity and interaction characteristics of the selected compounds with the target proteins. Both receptor and ligand entities were imported into PyRx and converted to the PDBQT format for docking. Site-targeted molecular docking was performed using AutoDock Vina in PyRx. A consistent Vina search approach was implemented for each receptor using the predetermined grid centers and box dimensions supplied in the configuration files. The docking grid was established at coordinates (x = 17.171, y = 25.151, z = 58.717) and had dimensions of (size_x = 23.269, size_y = 25.988, size_z = 38.277 Å) for 6UY4. The comprehensiveness was determined to be 8. The docking procedure for 4I07 was executed with the same exhaustiveness factor (8) and a grid positioned at (x = 1.160, y = 0.158, z = 1.786), employing box dimensions of (size_x = 61.816, size_y = 39.892, size_z = 53.323 Å). Replicate docking runs were performed using the same docking methodology across several computer systems to assess the reproducibility of the molecular docking data. The resultant docked complexes were then analysed for ligand-protein interactions and binding orientation inside the active-site pocket.

### 2.6. Docking validation using DUD-E decoy analysis

The molecular docking protocol was validated by retrospective virtual screening using the three highest-scoring active ligands and their corresponding directory of useful decoys, enhanced (DUD-E) benchmark datasets retrieved from the DUD-E server (https://dude.docking.org/). DUD-E contains experimentally validated active ligands from the ChEMBL database (https://www.ebi.ac.uk/chembl/) and property-matched decoy compounds from the ZINC database (https://zinc.docking.org/substances/home/), providing a robust framework for assessing docking performance. The active-decoy library was docked into the validated binding site of the target protein using the same docking parameters and scoring function as those used in the primary virtual screening, ensuring methodological consistency [31]. The docking scores were ranked based on predicted binding affinity, and a Receiver Operating Characteristic (ROC) curve was generated by plotting the true positive rate (TPR) against the false positive rate (FPR) across all classification thresholds. The area under the ROC curve (AUC) was determined to assess the docking protocol’s performance in discriminating active ligands from decoys, with values of 1.0 and 0.5 representing perfect and random performance, respectively. An AUC ≥ 0.70 was considered satisfactory predictive performance for virtual screening [32].

### 2.7. MD simulation

To understand the atomic stability and binding kinetics of the ligand-receptor complexes, MD simulations were performed using YASARA Structure (v. 25.12.1.W.64) with the AMBER14 force field, yielding 401 snapshots. This approach allows identification of conformational changes that occur over time and provides insight into the thermodynamic favorability of the interactions [33–35]. A cubic cell was used to solvate the compounds using the TIP3P water model at 0.997 g/cm3. The simulations were performed under physiological conditions at pH 7.4, 1 bar, and 298 K. To maintain realistic ion concentrations and electrical neutrality in the systems, 0.9% NaCl (Na and Cl counterions) was added. Initial steric conflicts were addressed through a rigorous minimisation procedure using the steepest-descent method, which resulted in the systems reaching a local energy minimum. All mobile atoms were integrated numerically using the equations of motion with periodic boundary conditions, with a time step of 2.5 fs for the complex and 1.25 fs for the complex. A combination of computational efficiency and actual physical accuracy was used in electrostatic management. Short-range van der Waals and Coulombic interactions were treated as cut-off interactions at 8 Å, and long-range electrostatic forces were treated as PME. After 100 ns of trajectories, many biophysical parameters were used to quantify the complexes’ structural stability and compactness [36] such as Root Mean Square Deviation (RMSD) of proteins and ligands, and Root Mean Square Fluctuation (RMSF) for structural equilibrium and residue-level flexibility, Radius of Gyration (Rg) for protein folding and compactness, Solvent Accessible Surface Area (SASA) for hydrophobic core changes, and Total Interaction Energy for binding interface strength and persistence.

### 2.8. Density functional theory (DFT) analysis

Selected ligands were initially chemically configured from the online structural database, PubChem (https://pubchem.ncbi.nlm.nih.gov/). Later, all quantum chemical calculations were performed using the Gaussian 09W (Revision D.01) software [37] within the DFT [38] at the B3- LYP level of theory [39] using a 6-31G basis set [40]. Highest Occupied Molecular Orbitals (HOMOs) have the potential to donate electrons to the protein, while the Lowest Unoccupied Molecular Orbitals (LUMOs) have the potential to accept electrons from the protein. The energy gap (ΔE), hardness (η), softness (S) and potentiality (μ) determinations were done using the following equations:

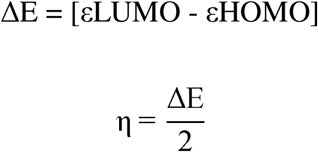

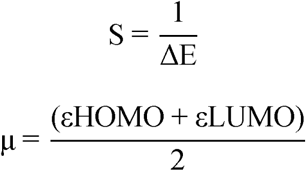

Based on the above determined equations, the higher the hardness, the lower the reactivity; the lower the hardness, the higher the reactivity [41]. The selected compounds were also analysed for the electrostatic potential (ESP) map.

### 2.9. QSAR analysis

Quantitative Structure-Activity Relationship (QSAR) is a method of quantum chemistry that relates a compound’s biological activity to its molecular structure [42]. SMILES of selected compounds were acquired from the PubChem database, and QSAR properties were calculated by the PASS online platform (http://way2drug.com/PassOnline/predict.phpp). The PASS algorithm uses a comprehensive database of over 4000 attributes to accurately recognize physiologically active compounds in the database with increased prediction reliability, including drug- and non-drug-related activities. The relevant prognostic variables are expressed as probabilities of activity (Pa) and inactivity (Pi), each ranging from 0.00 to 1.00. If the molecule is classified as neither completely inactive nor active at the same time, the criteria applied must be that Pa > Pi, which is to say the sum of Pa and Pi = 1 [43].

## 3. Results

### 3.1. Ligand selection and ADMET prediction

The ADMET prediction and drug-likeness screening were performed using SwissADME and the pkCSM server to identify fungal secondary metabolites with acceptable pharmacokinetic and toxicological profiles. As shown in **Table 1**, the initial dataset contained 1831 fungal secondary metabolites, which were reduced to 1812 unique compounds after duplicate removal. Following compound standardization based on compound name, identifiers, PubChem CID matching, and availability of 3D structures, 1647 compounds were retained for ADME screening. SwissADME-based drug-likeness filtering was then applied using criteria such as high gastrointestinal absorption, no blood-brain barrier permeability, no violations of Lipinski rules, zero PAINS alerts, and molecular weight below 500 Da. This filtering step reduced the compound pool to 266 compounds (**Table 1**). Subsequently, toxicity prediction was performed using the pkCSM server, where compounds were excluded if they showed predicted AMES toxicity, skin sensitization, hepatotoxicity, or hERG inhibition. After toxicity screening, 120 compounds passed all selection criteria and were finally selected for molecular docking analysis (**Table 1**). The full ADMET profiles of the final selected 120 compounds are provided in **Supplementary File 1**.

**Table 1.** Compound processing and selection criteria for docking analyses.

| Phases | Screening steps | Objectives | Criteria applied | Remaining compounds (n) |
| --- | --- | --- | --- | --- |
| 1 | Initial Dataset | Compile all collected fungal secondary metabolites from the database | Raw dataset | 1831 |
| 2 | De-duplication | Retain only unique chemical entities | Duplicate compounds removed | 1812 |
| 3 | Standardization | Standardize compound names, identifiers and structures | PubChem CID and exact compound name matching and available 3D structures | 1647 |
| 5 | ADME Filtering | Select compounds with favorable pharmacokinetic properties | High GI absorption; No BBB permeability; No Lipinski violations; Pains alert = 0; Molecular weight < 500 Da | 266 |
| 6 | Toxicity Filtering | Eliminate compounds with predicted toxicological risk | AMES = No; No skin sensitization; Hepatotoxicity = No; hERG inhibition = No | 120 |
| 7 | Final Selection | Identify compounds suitable for molecular docking analysis | Compounds passing all previous filters | 120 |

### 3.2. Molecular docking and validation

After removing duplicate entries and excluding ligands without available 3D conformers, 122 ligands were docked against four targets (6UY4 and 4I07). For 6UY4, the top three ligands with the best binding affinity were CIDs: 122184973, 132962217, and 139584993. In the case of 4I07, the top three ligands were CIDs: 122184973, 132962217 and 132962216. Notably, CID_122184973 and CID_132962217 consistently exhibited strong binding affinities toward both targets, indicating their potential as dual-target inhibitors against schistosomiasis (**Table 2**). The complete docking results for both 6UY4 and 4I07 are provided in **Supplementary File 2**.

**Table 2.** Top-ranked ligand compounds and predicted binding affinities against the target proteins 6UY4 and 4I07.

| Targets (PDB IDs) | Rank | Compound's names | Ligands (CIDs) | Binding affinities (kcal/mol) |
| --- | --- | --- | --- | --- |
| 6UY4 | 1 | Ganoderlactone B | 122184973 | -10.6 ± 0.06 |
|  | 2 | Spiroapplanatumine F | 132962217 | -10.0 ± 0.1 |
|  | 3 | Ganodernoid C | 139584993 | -9.9 ± 0.06 |
| 4I07 | 1 | Ganoderlactone B | 122184973 | -9.0 ± 0.12 |
|  | 2 | Spiroapplanatumine F | 132962217 | -8.9 ± 0.06 |
|  | 3 | Spiroapplanatumine E | 132962216 | -8.9 ± 0.06 |

The validity and discriminatory performance of the adopted docking protocol were evaluated using the three highest-ranked active compounds identified for each of the two target proteins (6UY4 and 4I07). Following the Directory of Useful Decoys, enhanced (DUD-E) recommendation of 50 decoys per active ligand, 150 physicochemically matched decoy molecules were generated for each target, resulting in a validation library comprising 153 compounds (3 active ligands and 150 decoys). The discriminatory performance of the docking protocol was assessed by constructing ROC curves, in which TPR was plotted against FPR across all classification thresholds (**Fig 1**). The resulting area under the ROC curve (AUC) values were 0.98 for 6UY4 and 0.96 for 4I07, demonstrating excellent enrichment and discriminatory power. These AUC values indicate that the docking protocol correctly ranks an active ligand above a randomly selected decoy with probabilities of approximately 98% and 96% for 6UY4 and 4I07, respectively. Both values are substantially higher than the random classification threshold (AUC = 0.5) and exceed the commonly accepted benchmark for satisfactory predictive performance (AUC ≥ 0.70) in virtual screening, confirming the robustness and statistical validity of the adopted docking protocol in distinguishing active ligands from inactive decoys.

**Fig 1.**
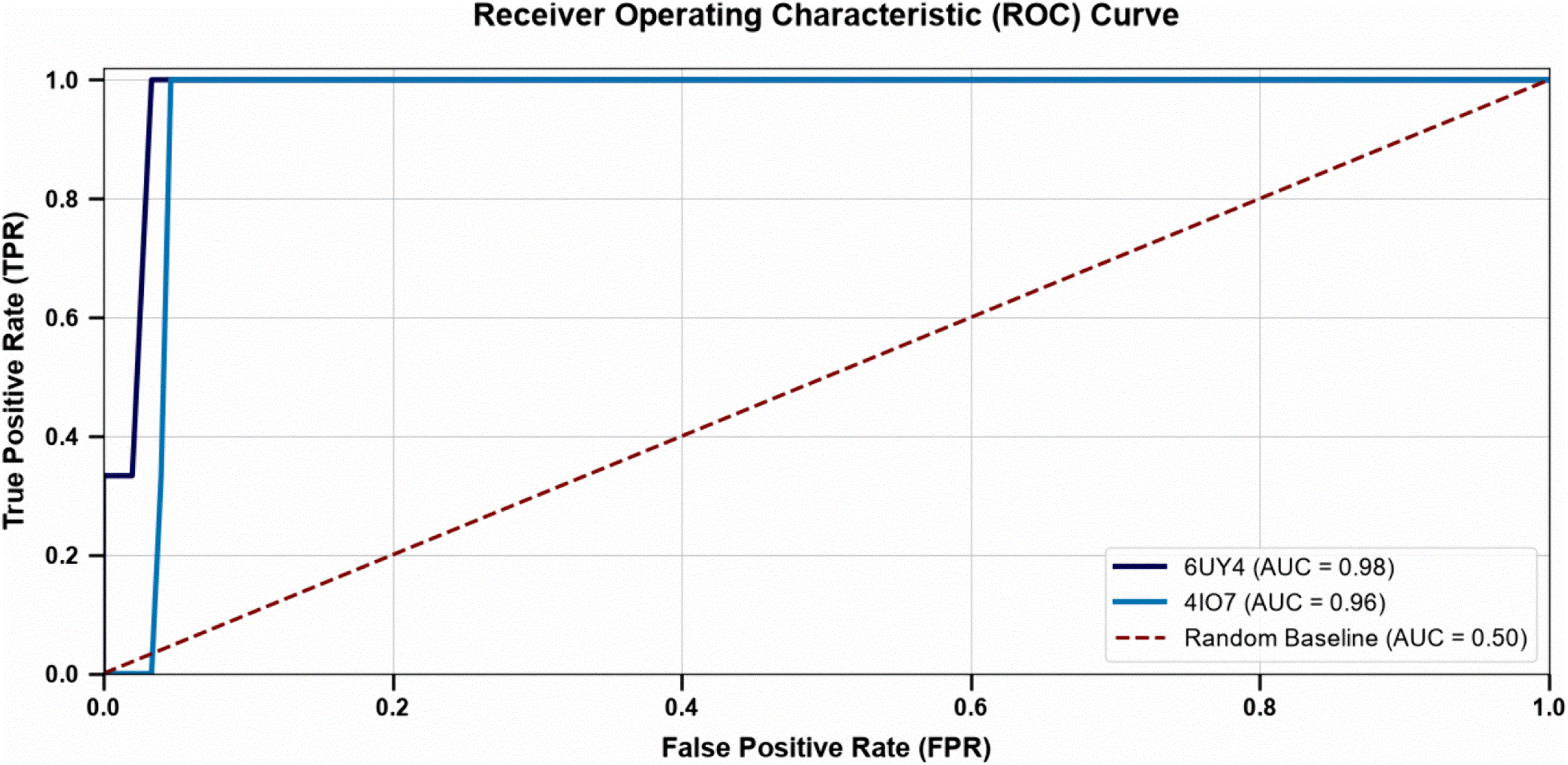
Receiver Operating Characteristic (ROC) curves evaluating the discriminatory performance of the docking protocol.

ROC curves showing the validation of the molecular docking protocol for target proteins 6UY4 and 4I07 using DUD-E active-decoy datasets. The docking protocol demonstrated excellent discriminatory performance, with AUC values of 0.98 for 6UY4 and 0.96 for 4I07. The dashed red diagonal line represents the performance of a random classifier (AUC = 0.50).

### 3.3. Evaluation of Protein-ligand Interactions

The top three docked compounds (CIDs: 122184973, 132962217, and 139584993) with 6UY4 (**Fig 2**) and the other top three compounds (CIDs: 122184973, 132962216, 132962217) in complex with 4I07 (**Fig 3**) were selected for further downstream research, based on their binding affinity. For this purpose, Discovery Studio 2021 Client was utilised to visualise molecular interactions. The protein-ligand interaction profiles of the selected compounds against the 6UY4 and 4I07 targets were further investigated using both two- and three-dimensional interaction mapping. The major intermolecular contacts, including conventional hydrogen bonds, π interactions, and van der Waals contacts, were identified to determine which residues contribute to ligand stabilization within the binding pocket. The interaction patterns revealed that only a subset of residues consistently participated in strong binding, indicating the presence of key hotspot residues within each target site. The detailed interaction profiles for each PubChem compound are summarized in **Table 3**.

**Fig 2.**
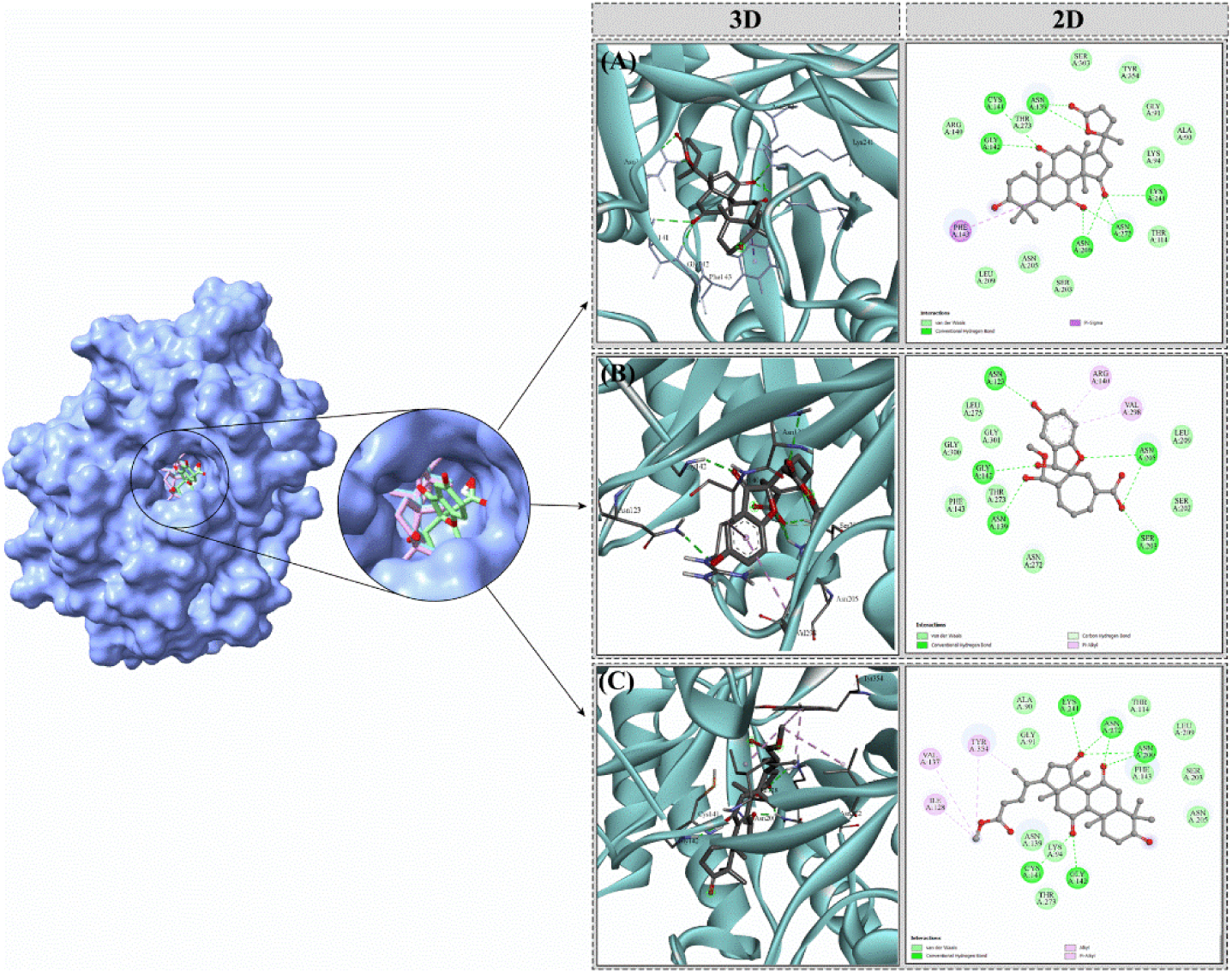
Comparative 3D binding conformations and 2D interaction maps of CID_122184973 (A), CID_132962217 (B), and CID_139584993 (C) within the active site of 6UY4.

**Fig 3.**
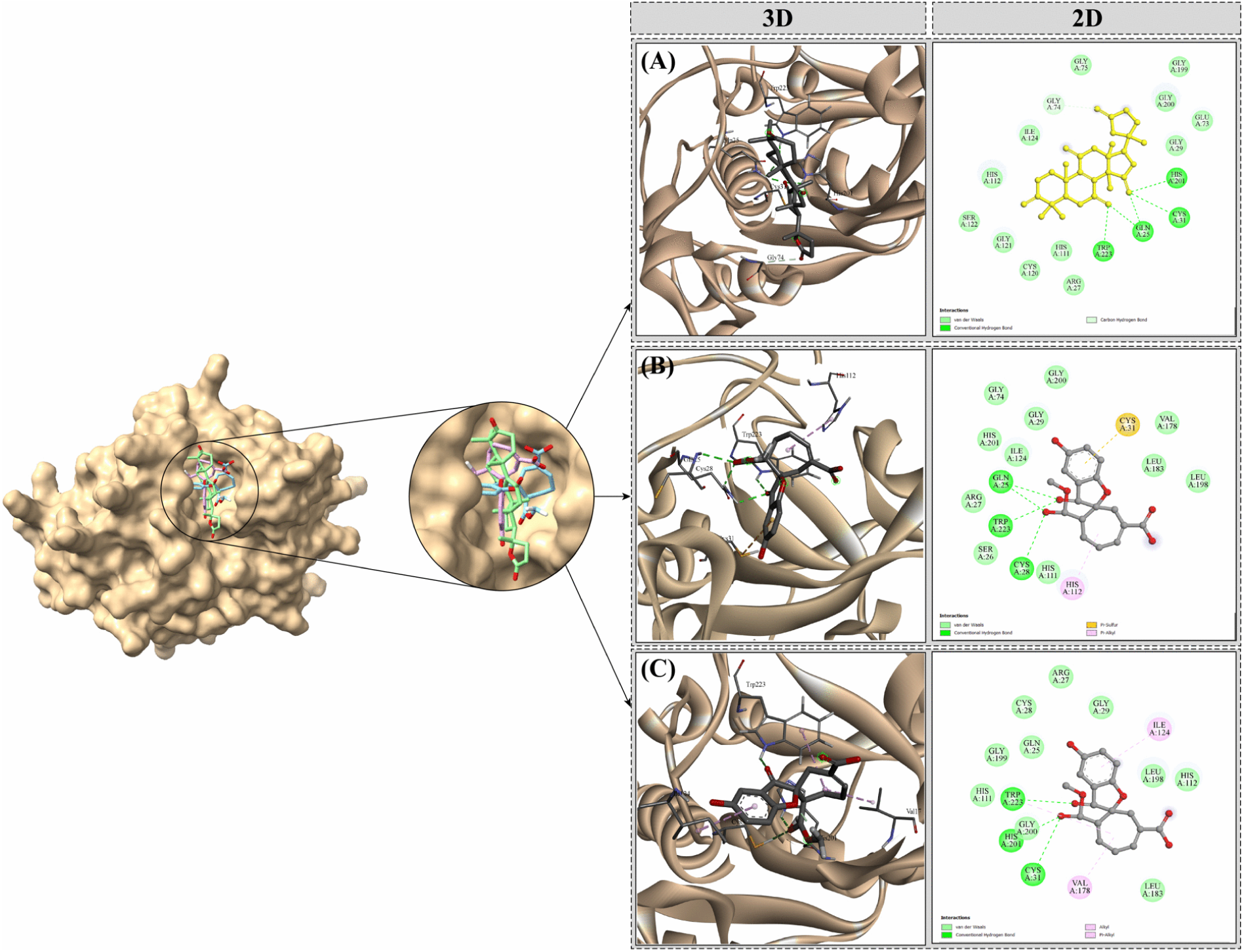
Comparative 3D binding poses and 2D interaction profiles of CID_122184973 (A), CID_132962216 (B), and CID_132962217 (C) within the active site of 4I07.

**Table 3.** Comparative protein-ligand interaction profile of selected PubChem compounds with 6UY4 and 4I07 protein-binding pockets.

| Proteins | PubChem CIDs | Hydrogen Bonds | π-Interactions | Van Der Waals Interactions | Others |
| --- | --- | --- | --- | --- | --- |
| 6UY4 | CID_122184973 | ASN139,<br>GLY142, LYS241,<br>ASN272,<br>ASN200, CYS141 | PHE143 | ARG140, ALA90, GLY91, LYS94,<br>SER203, ASN205, SER303,<br>TYR354, THR273 | - |
|  | CID_132962217 | ASN123,<br>GLY142,<br>ASN139,<br>ASN205, SER203 | ARG140,<br>VAL298 | LEU275, GLY300, GLY301,<br>THR273, PHE143, ASN272,<br>LEU209, SER202 | - |
|  | CID_139584993 | LYS241, ASN272,<br>ASN200,<br>GLY142, CYS141 | TYR354 | ALA90, GLY91, LYS94, THR114,<br>ASN139, THR273, PHE143,<br>SER203, ASN205, LEU209 | VAL135,<br>ILE128 |
| 4I07 | CID_122184973 | HIS201, GLN25,<br>TRP223, CYS31,<br>GLY74 | - | GLY75, GLY119, GLY200, GLU73,<br>GLY29, HIS112, SER122, GLY121,<br>HIS111, ARG27, ILE124 | - |
|  | CID_132962216 | GLN25, TRP223,<br>CYS28 | HIS112,<br>CYS31 | GLY74, GLY200, GLY29, HIS201,<br>ILE124, ARG27, SER26, HIS111,<br>VAL178, LEU183, LEU198 | - |
|  | CID_132962217 | TRP223, HIS201,<br>CYS31 | VAL178 | ARG27, CYS28, GLY29, GLY199,<br>GLY200, HIS112, LEU183, LEU198 | ILE124 |

### 3.4. MD simulations

#### 3.4.1. Protein RMSD analysis

The stability of the *S. mansoni* survival proteins 4I07 and 6UY4 in complex with selected fungal metabolites was assessed via molecular dynamics simulations over 100 ns, and protein backbone RMSD values were analyzed. For 4I07, all three ligand complexes (CID_122184973, CID_132962216, CID_132962217) exhibited initial fluctuations within the first 5 ns, after which RMSD values stabilized around 1.2-1.6 Å, indicating minor conformational adjustments and overall structural stability throughout the simulation (**Fig 4a**). Among these, CID_132962216 showed slightly higher deviations, reaching up to ∼1.7 Å, while CID_122184973 and CID_132962217 remained below 1.5 Å on average. Similarly, 6UY4 complexes with CID_122184973, CID_132962217, and CID_139584993 displayed initial increases in RMSD during the first 10 ns, followed by stabilization between 1.4 and 1.7 Å (**Fig 4b**). Notably, CID_132962217 maintained a consistent RMSD across both proteins, suggesting a particularly stable interaction. The RMSD analysis confirms that the common ligands CID_122184973 and CID_132962217 form stable complexes with both 4I07 and 6UY4, supporting their potential as lead candidates for further investigation.

**Fig 4.**
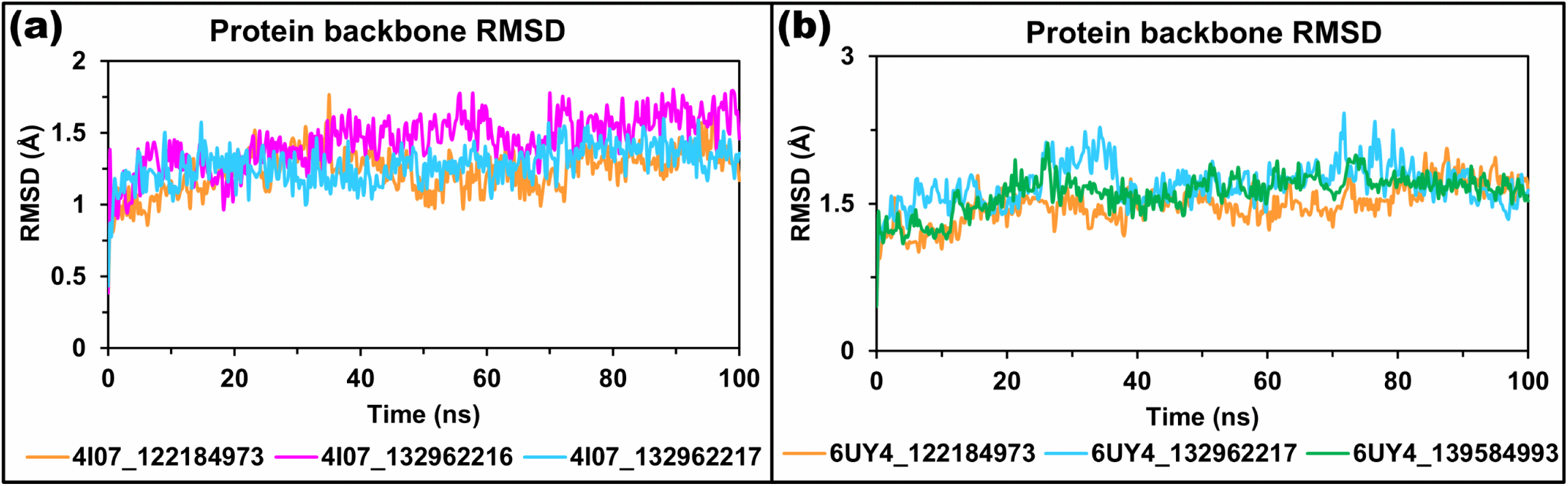
Backbone RMSD analysis of *S. mansoni* survival proteins 4I07 (a) and 6UY4 (b) in complex with selected fungal metabolites over 100 ns of MD simulation. RMSD values (Å) were monitored for **(a**) 4I07 complexes with CID_122184973 (orange), CID_132962216 (magenta), and CID_132962217 (cyan), and for **(b)** 6UY4 complexes with CID_122184973 (orange), CID_132962217 (cyan), and CID_139584993 (green).

#### 3.4.2. Ligand RMSD analysis

Ligand RMSD analysis was conducted to assess the dynamic behavior and stability of the fungal metabolites within the binding pockets. For 4I07 (**Fig 5a**), CID_132962217 displayed consistently low RMSD values (∼1-5 Å) throughout the simulation, indicating that it remained well-anchored within the binding site with minimal structural displacement. In contrast, CID_122184973 and CID_132962216 exhibited higher fluctuations, with RMSD values increasing up to 60-70 Å, suggesting that these ligands experienced significant repositioning or partial egress from the binding pocket, likely reflecting weaker or more flexible interactions. For 6UY4 (**Fig 5b**), all three ligands (CID_122184973, CID_132962217, and CID_139584993) showed lower and more uniform RMSD values (∼1.5-6 Å), indicating that the ligands maintained stable conformations and strong interactions within the binding site. Importantly, the two ligands shared by both proteins, CID_122184973 and CID_132962217, exhibited consistently low and stable RMSDs across both simulations, reflecting their ability to form robust, persistent interactions with the target proteins. These findings suggest that these two compounds are not only well accommodated within the binding sites but also structurally restrained, reinforcing their potential as lead compounds for further *in silico* and experimental validation against *S. mansoni* survival proteins.

**Fig 5.**
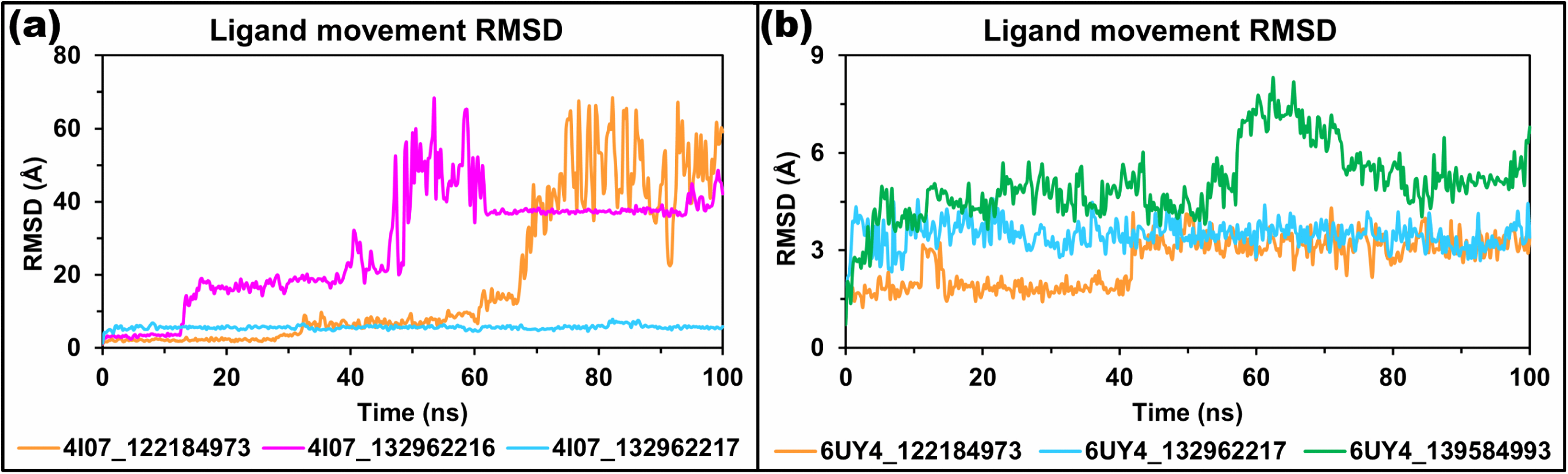
Ligand RMSD analysis of fungal metabolites bound to *Schistosoma mansoni* proteins. (a) 4I07 and (b) 6UY4 over 100 ns of MD simulation.

#### 3.4.3. RMSF analysis

RMSF measures the average positional fluctuation of each residue during the simulation and is used to identify flexible and stable regions of a protein-ligand complex. Lower RMSF values indicate restricted residue motion and structural stability, whereas higher RMSF values indicate increased local flexibility, commonly observed in terminal or loop regions [44]. In the 4I07 complexes, residue-wise RMSF values remained generally low, indicating stable protein conformations during simulation. The mean RMSF values were 1.18 Å for 4I07_122184973, 1.13 Å for 4I07_132962216, and 0.96 Å for 4I07_132962217, showing that 4I07_132962217 had the most rigid residue profile. The overall average RMSF across the three 4I07 systems was 1.09 Å, with only 17 of 254 residues showing average RMSF values above 2.0 Å. The highest fluctuation was observed at the N-terminal VAL1 residue, with RMSF values of 4.54, 4.53, and 4.66 Å, respectively. Other flexible residues included LYS114 (2.82 Å), GLU2 (2.63 Å), ASN53 (2.57 Å), ARG129 (2.53 Å), and LYS139 (2.46 Å). In contrast, regions such as residues 38-46, 88-92, 100-106, 169-175, and 220-224 showed average RMSF values below 0.8 Å, reflecting stable structural segments (**Fig 6a**). For 6UY4, the RMSF profiles showed slightly greater flexibility than those for 4I07. The mean RMSF values were 1.20 Å for 6UY4_122184973, 1.19 Å for 6UY4_132962217, and 1.31 Å for 6UY4_139584993, indicating that 6UY4_139584993 had the greatest residue mobility. The overall average RMSF was 1.23 Å, and 26 of 354 residues showed average RMSF values above 2.0 Å. The highest fluctuation occurred at the N-terminal SER25, with RMSF values of 5.85, 8.58, and 5.25 Å, respectively, and an average RMSF of 6.56 Å. Other highly flexible residues were GLY26 (4.65 Å), ASN27 (4.32 Å), GLU28 (3.29 Å), GLN295 (4.44 Å), MET377 (4.32 Å), and THR378 (5.46 Å). Stable regions with average RMSF values below 1.0 Å were observed at residues 82-91, 105-117, 135-152, 164-171, 194-198, 269- 273, 299-304, 327-334, 346-350, and 358-363 (**Fig 6b**). Overall, both proteins showed stable residue behavior, with flexibility mainly limited to terminal and loop regions rather than widespread structural destabilization.

**Fig 6.**
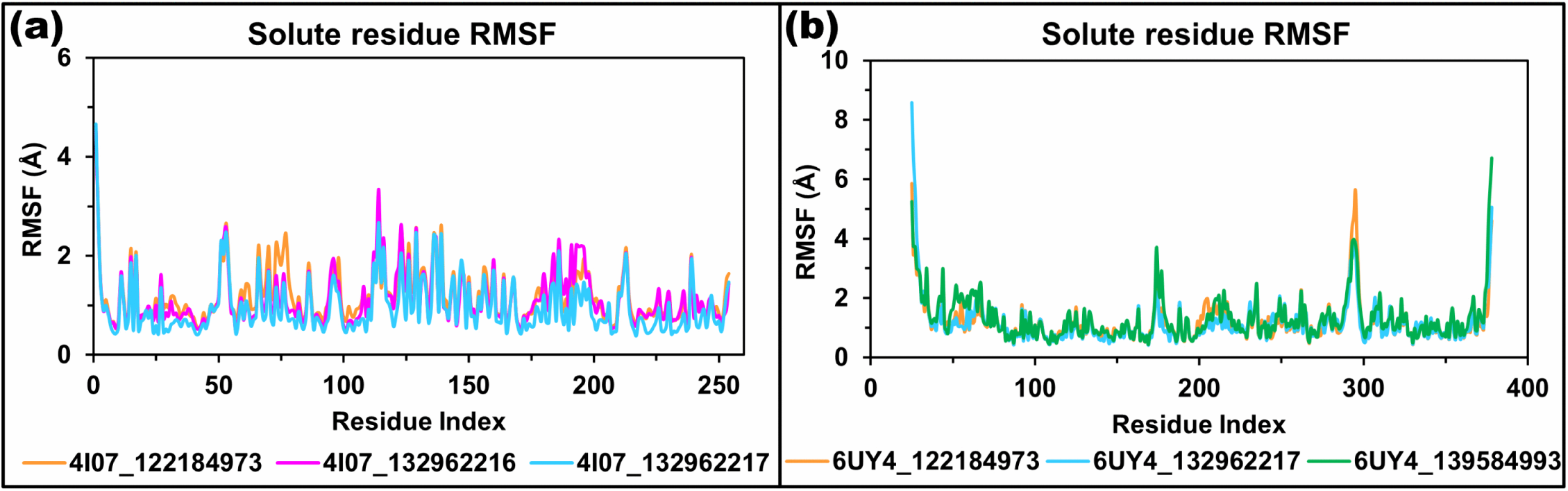
Residue-wise RMSF analysis of simulated protein-ligand complexes. **(a)** RMSF plot of 4I07 in complex with CID_122184973, CID_132962216, and CID_132962217, and **(b)** 6UY4 in complex with CID_122184973, CID_132962217, and CID_139584993.

#### 3.4.4. Rg analysis

Rg measures the overall compactness of a protein structure around its center of mass. In MD simulation, Rg analysis is used to assess whether the protein-ligand complex remains structurally compact and stable or undergoes expansion/unfolding during the simulation period. In the 4I07 complexes, Rg values showed ligand-dependent stability. The 4I07_132962217 complex maintained the most compact and stable profile, with an average Rg of 17.762 ± 0.060 Å and a narrow range of 17.574-17.986 Å throughout 100 ns. The 4I07_132962216 complex also remained relatively stable, although a temporary rise was observed around 50-60 ns, reaching 19.074 Å at 58.75 ns, before returning to a compact state. Its average Rg was 18.086 ± 0.202 Å. In contrast, 4I07_122184973 showed greater fluctuation after ∼70 ns, with a maximum Rg of 19.379 Å at 78.50 ns and an average value of 17.981 ± 0.419 Å, indicating comparatively higher conformational expansion (**Fig 7a**). For the 6UY4 complexes, all systems maintained stable Rg profiles within a narrow range, suggesting good structural compactness during the simulation. The average Rg values were 19.742 ± 0.113 Å for 6UY4_122184973, 19.805 ± 0.086 Å for 6UY4_132962217, and 19.727 ± 0.088 Å for 6UY4_139584993. Although 6UY4_132962217 showed a small transient peak of 20.170 Å at 71.75 ns, it remained stable overall (**Fig 7b**). Therefore, the Rg analysis indicates that CID_132962217 provided the highest compactness in the 4I07 system, while all three 6UY4 complexes preserved stable structural compactness throughout the simulation.

**Fig 7.**
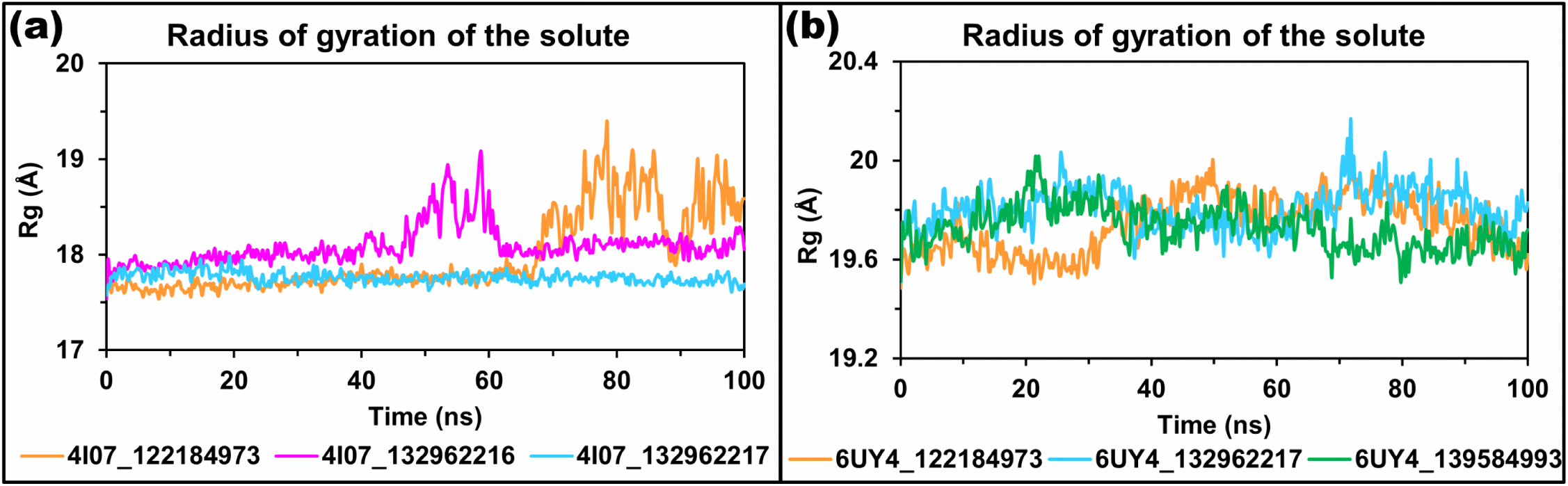
Radius of gyration (Rg) analysis of protein-ligand complexes during 100 ns MD simulation. **(a)** Rg profiles of 4I07 complexed with CID_122184973, CID_132962216, and CID_132962217. The 4I07_132962217 complex maintained the most stable and compact Rg pattern, whereas 4I07_122184973 showed increased fluctuation after ∼70 ns. **(b)** Rg profiles of 6UY4 complexed with CID_122184973, CID_132962217, and CID_139584993. All 6UY4 complexes showed narrow Rg fluctuations, indicating stable structural compactness throughout the simulation.

#### 3.4.5. SASA analysis

SASA measures the total surface area of a protein that is exposed to the surrounding solvent. In MD simulations, SASA analysis is used to evaluate changes in protein surface exposure, compactness, and possible conformational rearrangement after ligand binding; a stable SASA profile generally indicates preservation of structural stability. In the 4I07 complexes, SASA values showed ligand-specific variation during the 100 ns simulation. The 4I07_132962217 complex exhibited the lowest average SASA (11901.758 ± 163.215 Å^2^) with a comparatively narrow range of 11427.549-12315.392 Å^2^, indicating better maintenance of a compact and less solvent-exposed structure. The 4I07_132962216 complex showed a higher average SASA (12303.659 ± 246.879 Å^2^) and reached a maximum of 13118.971 Å^2^ at 58.75 ns, suggesting a transient increase in solvent exposure. The 4I07_122184973 complex also showed increased fluctuation, especially after ∼65 ns, with an average SASA of 12098.275 ± 370.840 Å^2^ and a maximum of 12955.314 Å^2^ at 81.75 ns. Overall, CID_132962217 produced the most stable SASA profile for 4I07 (**Fig 8a**). For the 6UY4 complexes, SASA values remained relatively stable, although moderate fluctuations were observed among the ligand-bound systems. The 6UY4_139584993 complex showed the lowest average SASA (15521.716 ± 291.189 Å^2^), suggesting comparatively lower solvent exposure. The 6UY4_122184973 complex maintained an average SASA of 15608.255 ± 323.462 Å^2^, with a maximum of 16355.348 Å^2^ at 75.25 ns. The 6UY4_132962217 complex showed a slightly higher average SASA (15730.747 ± 240.194 Å^2^) and reached a transient peak of 16488.011 Å^2^ at 72.25 ns, but remained stable overall (**Fig 8b**). Therefore, the SASA analysis indicates that all complexes retained their structural integrity throughout the simulation, with 4I07_132962217 and 6UY4_139584993 showing comparatively lower solvent exposure.

**Fig 8.**
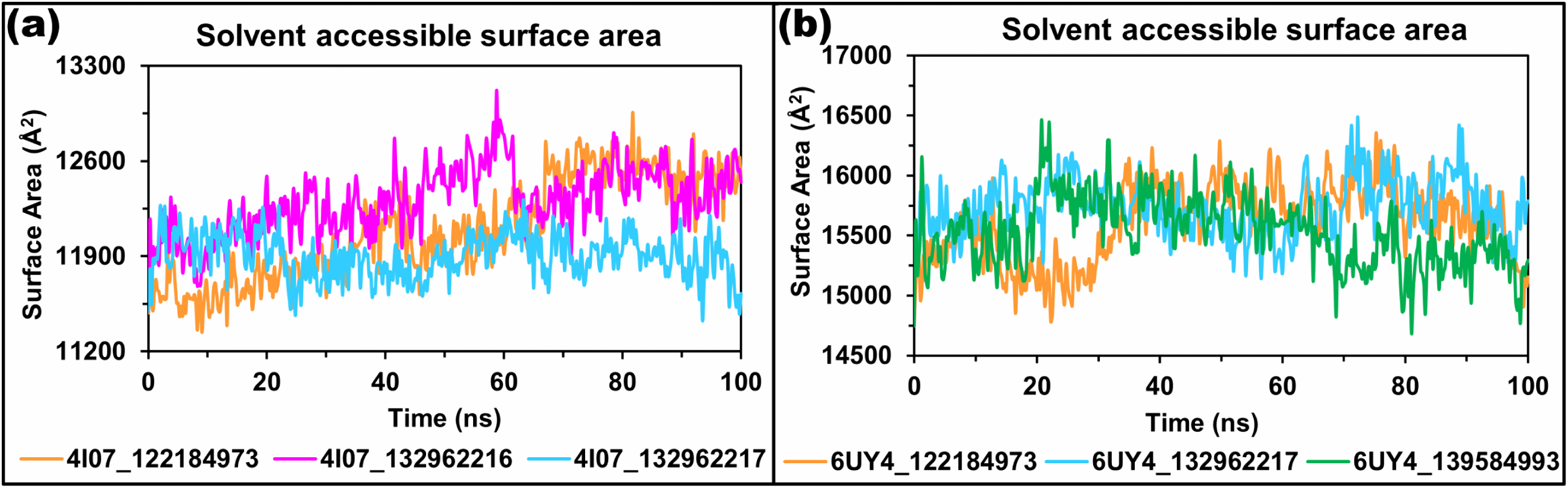
SASA analysis of protein-ligand complexes during 100 ns MD simulation. **(a)** SASA profiles of 4I07 complexes with CID_122184973, CID_132962216, and CID_132962217. **(b)** SASA profiles of 6UY4 complexes with CID_122184973, CID_132962217, and CID_139584993. Lower, stable SASA values indicate reduced solvent exposure and greater structural compactness. Overall, the selected complexes exhibited stable SASA patterns, indicating conformational stability during simulation.

#### 3.4.6. MM-PBSA analysis

The MM-PBSA binding energy analysis was performed using YASARA to evaluate the energetic stability of the selected fungal metabolites with the two *Schistosoma mansoni* survival proteins, 4I07 and 6UY4. According to the YASARA binding energy convention, higher or more positive binding energy values indicate more favorable binding [45]. For the 4I07 complexes, CID_122184973 showed the most favorable binding profile, with the highest average binding energy of −23.01 ± 43.01 kJ/mol. This ligand also reached a maximum binding energy of 117.37 kJ/mol, indicating a comparatively stronger interaction during several stages of the simulation. CID_132962216 and CID_132962217 showed lower average binding energies of −114.33 ± 85.51 kJ/mol and −115.27 ± 51.86 kJ/mol, respectively. Although CID_132962216 reached a positive maximum value of 54.69 kJ/mol, its high standard deviation indicates greater fluctuation throughout the trajectory. CID_132962217 displayed comparatively lower fluctuation than CID_132962216, but its average binding energy remained slightly less favorable than that of CID_132962216 and markedly lower than that of CID_122184973 (**Fig 9a**). Therefore, among the three ligands tested against 4I07, CID_122184973 demonstrated the most favorable YASARA-based MM-PBSA binding profile. For the 6UY4 complexes, CID_139584993 showed the most favorable binding energy, with the highest average value of -47.24 ± 34.56 kJ/mol, followed by CID_122184973 with −70.19 ± 36.05 kJ/mol. CID_139584993 also reached a maximum binding energy of 38.27 kJ/mol, suggesting a stronger, more favorable interaction during parts of the simulation. CID_122184973 showed a moderate binding profile and achieved a maximum value of 6.99 kJ/mol. In contrast, CID_132962217 showed the lowest average binding energy of −151.04 ± 37.53 kJ/mol, indicating the least favorable interaction with 6UY4 under the YASARA binding energy convention (**Fig 9b**). Overall, the MM-PBSA results suggest that CID_122184973 is the most promising common ligand against both 4I07 and 6UY4, while CID_139584993 showed the strongest binding profile specifically against 6UY4. Among the two common compounds, CID_122184973 performed better than CID_132962217 for both target proteins.

**Fig 9.**
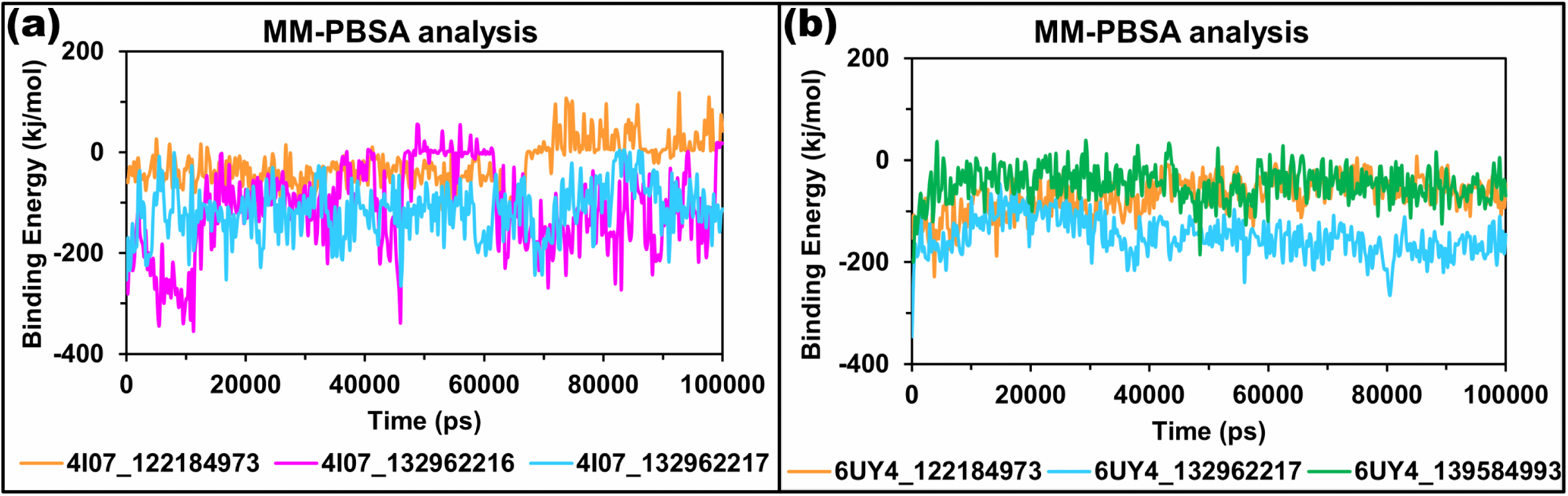
MM-PBSA binding energy analysis of selected fungal metabolites with *S. mansoni* survival proteins during a 100 ns simulation. The time-dependent MM-PBSA binding energy profiles were calculated from YASARA simulation trajectories for the docked protein-ligand complexes. **(a)** The left panel represents the binding energy fluctuations of the 4I07 complexed with CID_122184973, CID_132962216, and CID_132962217, while **(b)** the right panel represents the 6UY4 complexed with CID_122184973, CID_132962217, and CID_139584993.

#### 3.4.7. PCA analysis

Principal component analysis (PCA) was used to examine the dominant conformational motions of the simulated protein-ligand complexes in the PC1-PC2 subspace. For the 4I07 complexes, the first two principal components explained a substantial portion of the total motion, ranging from 71.23% to 78.70%. The 4I07_122184973 complex showed PC1 and PC2 contributions of 40.08% and 31.15%, respectively, with a broad distribution of conformations across PC1 values from 0.64 to 65.89 and PC2 values from −77.50 to −33.06. This wider spread suggests that the complex sampled a relatively broad conformational space during simulation. The 4I07_132962216 complex exhibited the highest cumulative variance among the 4I07 systems, with PC1 and PC2 explaining 54.17% and 24.53% of the motion variance, respectively. Its projection formed several separated clusters, indicating that the complex explored multiple conformational substates. In contrast, the 4I07_132962217 complex showed a more compact distribution, with PC1 and PC2 accounting for 47.65% and 29.77% of the variance, respectively (**Fig 10**). The coordinates were mainly concentrated within a narrower range, with the lowest dispersion among the 4I07 complexes. This pattern indicates more restricted conformational fluctuation and a comparatively stable conformational ensemble for the 4I07_132962217 complex.

**Fig 10.**
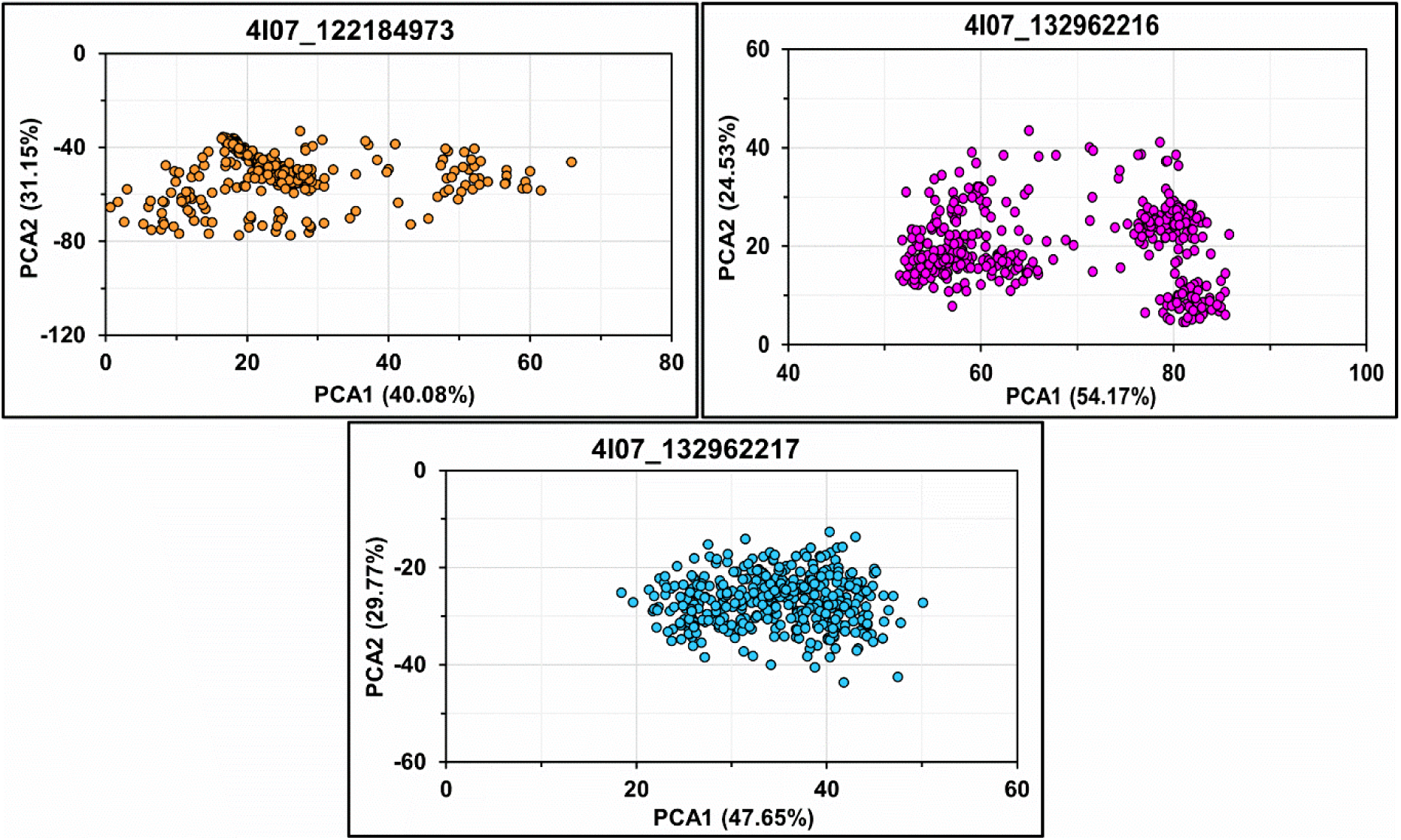
Principal component analysis of the 4I07 protein-ligand complexes. PCA scatter plots showing the projection of molecular dynamics trajectories onto the first two principal components for 4I07 complexed with three fungal metabolites: CID_122184973, CID_132962216, and CID_132962217.

For the 6UY4 complexes, PC1 and PC2 captured an even larger proportion of the total variance, ranging from 82.00% to 84.57%. The 6UY4-CID_122184973 complex showed PC1 and PC2 contributions of 47.93% and 36.64%, respectively, giving a cumulative variance of 84.57%. The conformations were distributed over PC1 values from 93.26 to 153.11 and PC2 values from 36.78 to 96.10, suggesting notable conformational sampling within the essential motion space. The 6UY4_132962217 complex showed the highest PC1 contribution among all systems, with PC1 explaining 58.27% and PC2 explaining 25.28% of the variance. Its PCA plot showed a wider, multi-clustered distribution, with PC1 values ranging from −108.46 to −39.42 and PC2 values from 12.86 to 68.37. This indicates that the complex underwent several conformational substates during the simulation. Similarly, the 6UY4_139584993 complex showed PC1 and PC2 contributions of 56.58% and 25.42%, respectively (**Fig 11**), with a cumulative variance of 82.00%. The projection formed distinct clusters across the PC1-PC2 space, reflecting broader conformational rearrangement rather than a single compact conformational basin.

**Fig 11.**
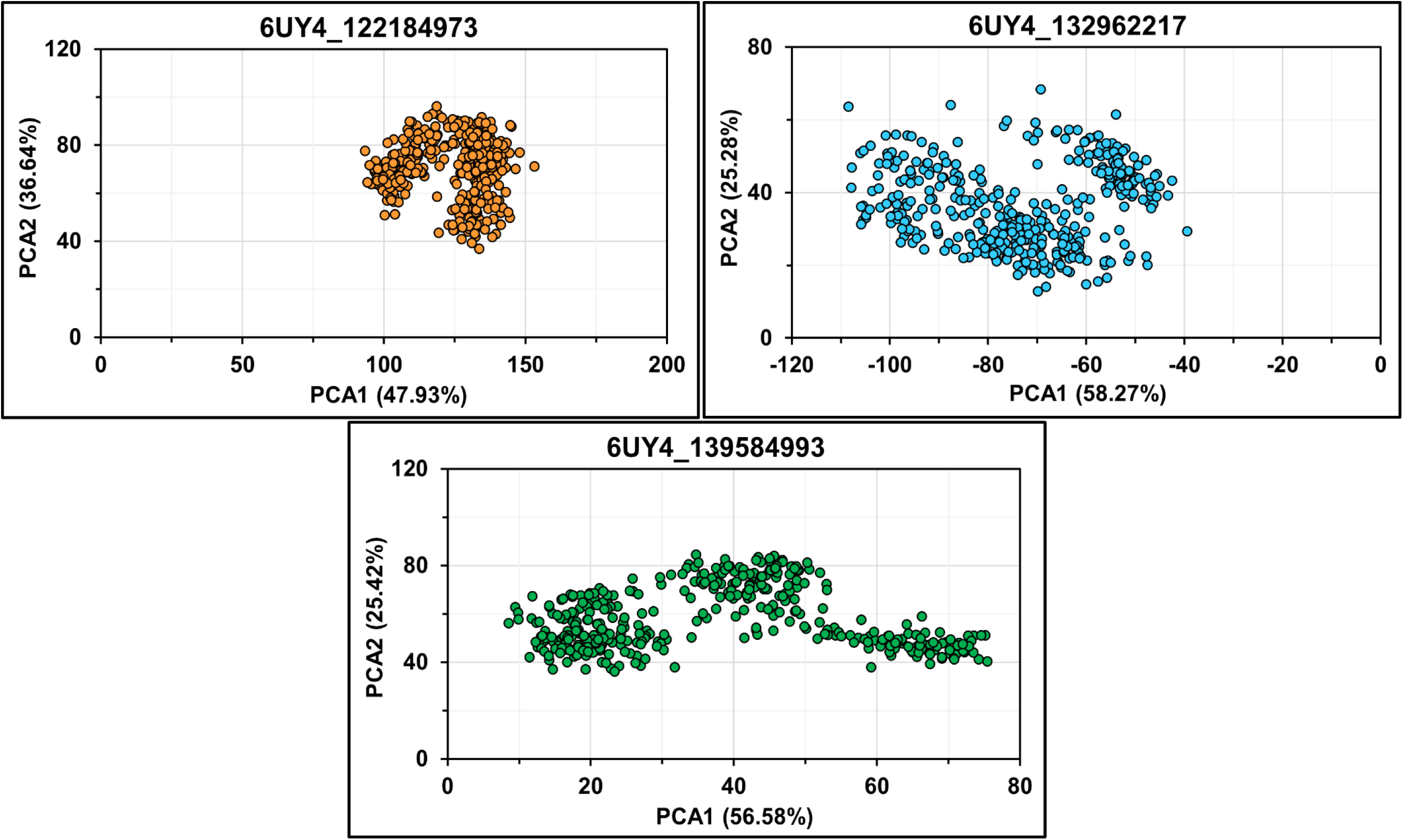
Principal component analysis of the 6UY4 protein-ligand complexes. PCA scatter plots showing the dominant conformational motions of 6UY4 in complex with CID_122184973, CID_132962217, and CID_139584993 during molecular dynamics simulation.

Overall, the PCA results indicate that the first two principal components captured the major collective motions of all simulated complexes. Among the common ligands, CID_122184973 exhibited broader conformational sampling for both proteins, particularly for 6UY4. CID_132962217 exhibited different behavior between the two targets: it produced a more compact, more restrained PCA distribution with 4I07, whereas it generated a wider, multi-basin conformational pattern with 6UY4. Therefore, based on the PC1-PC2 projections, CID_132962217 appears to stabilize the conformational space of 4I07 more effectively, whereas 6UY4 complexes generally retained broader conformational flexibility under the tested ligand-bound conditions.

### 3.5. DFT analysis

The frontier molecular orbital analysis revealed significant differences in the electronic behaviour of the selected compounds (CIDs: 122184973, 132962217, 139584993, 132962216) (**Fig 12**). The lowest HOMO-LUMO energy gap (3.552 eV) and hardness value (1.776 eV) were found for CID_139584993 among the investigated ligands, resulting in relatively high chemical reactivity and kinetic stability. CID_132962216 had the highest energy gap (4.133 eV) and hardness (2.0665 eV), indicating higher stability and lower reactivity. Similarly, the higher softness values for CID_139584993 and CID_122184973 suggest a greater tendency to interact via electron transfer. In summary, the detailed HOMO, LUMO and related global reactivity descriptors are provided in **Table 4**.

**Fig 12.**
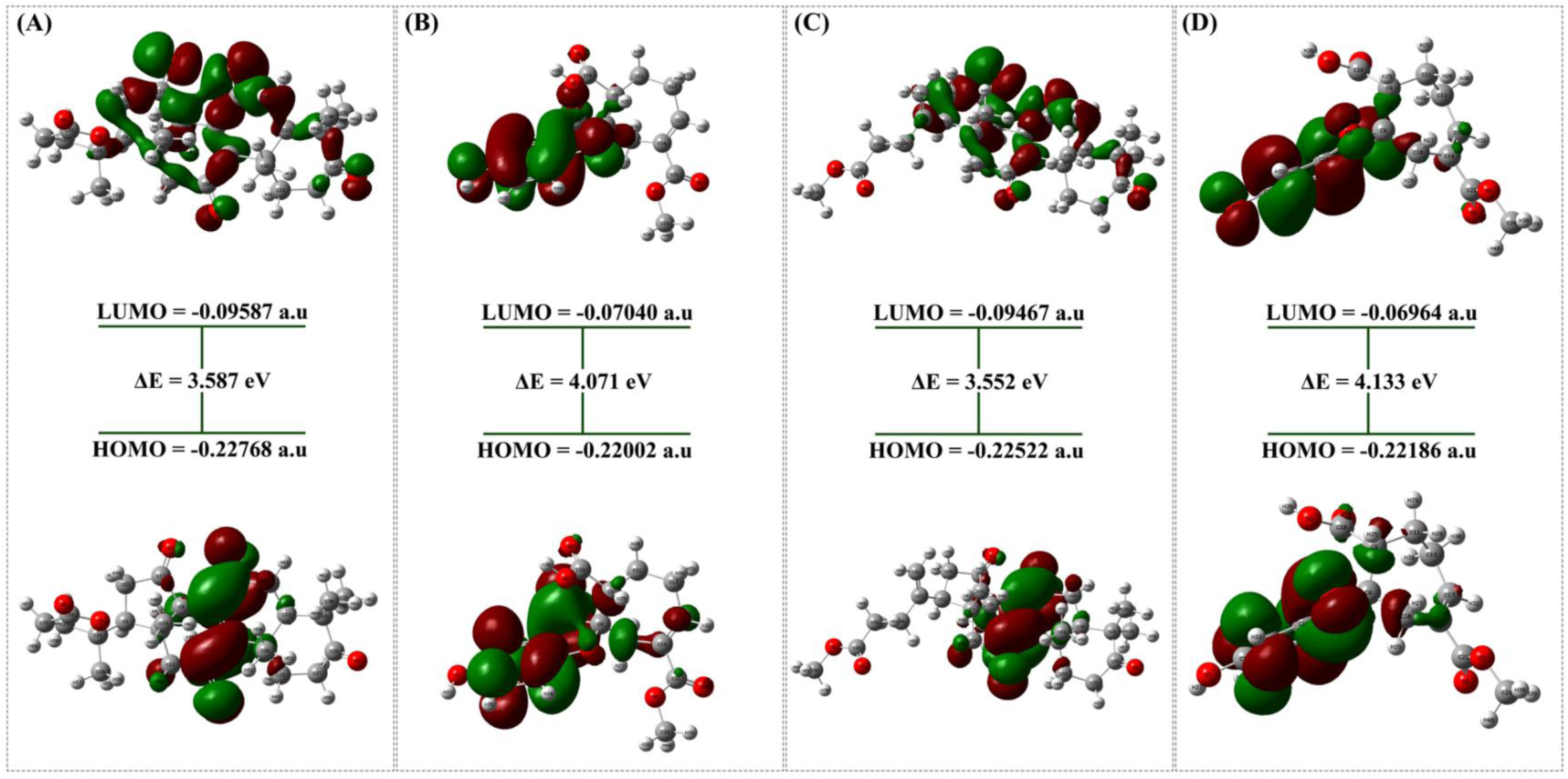
Frontier Molecular Orbital (HOMO-LUMO) distribution and energy gap analysis of the selected compounds: CIDs: 122184973 (A), 132962217 (B), 139584993 (C), and 132962216 (D).

**Table 4.** DFT-derived frontier orbital energies and associated global chemical reactivity descriptors of the investigated ligands (CIDs: 122184973, 132962217, 139584993, and 132962216).

| PubChem CIDs | $\epsilon_{\text{HOMO}}$ (au) | $\epsilon_{\text{LUMO}}$ (au) | $\Delta E$ (au) | $\Delta E$ (eV) | $\chi$ (eV) | $S$ (eV<sup>-1</sup>) | $\mu$ (eV) |
| --- | --- | --- | --- | --- | --- | --- | --- |
| CID_122184973 | -0.228 | -0.096 | 0.132 | 3.587 | 1.794 | 0.279 | -4.402 |
| CID_132962217 | -0.220 | -0.070 | 0.150 | 4.071 | 2.036 | 0.246 | -3.951 |
| CID_139584993 | -0.225 | -0.095 | 0.131 | 3.552 | 1.776 | 0.282 | -4.353 |
| CID_132962216 | -0.222 | -0.070 | 0.152 | 4.133 | 2.067 | 0.242 | -3.966 |

### 3.6. ESP Calculation

The ESP map is an important illustration in molecular recognition, which is employed to compare the relative polarity of a molecule [46]. **Fig 13** illustrates the ESP maps of the chosen compounds (CIDs: 122184973, 132962217, 139584993, and 132962216). The coloration of the map represents different electrostatic potential zones. The ESP map of the selected compounds showed that the oxygen atoms exhibited the most negative ESP, while the positive ESP was the highest for the hydrogen atoms. The positive electrostatic potential spectrum ranged from +7.493 to +8.054 au, and the negative electrostatic potential ranged from −7.493 to -8.054 au. It is important to note that CID_139584993 also has the highest positive potential (+8.054 au) and the highest negative potential (-8.054 au). CID_132962216, on the other hand, has the lowest positive potential at +7.493 au and the lowest negative potential at −7.493 au. In addition, CIDs: 122184973 and 132962217 manifested moderate positive electrostatic potentials of (+7.530 au) and (+8.035 au), respectively, as well as negative electrostatic potentials of (-7.530 au) and (- 8.035 au).

**Fig 13.**
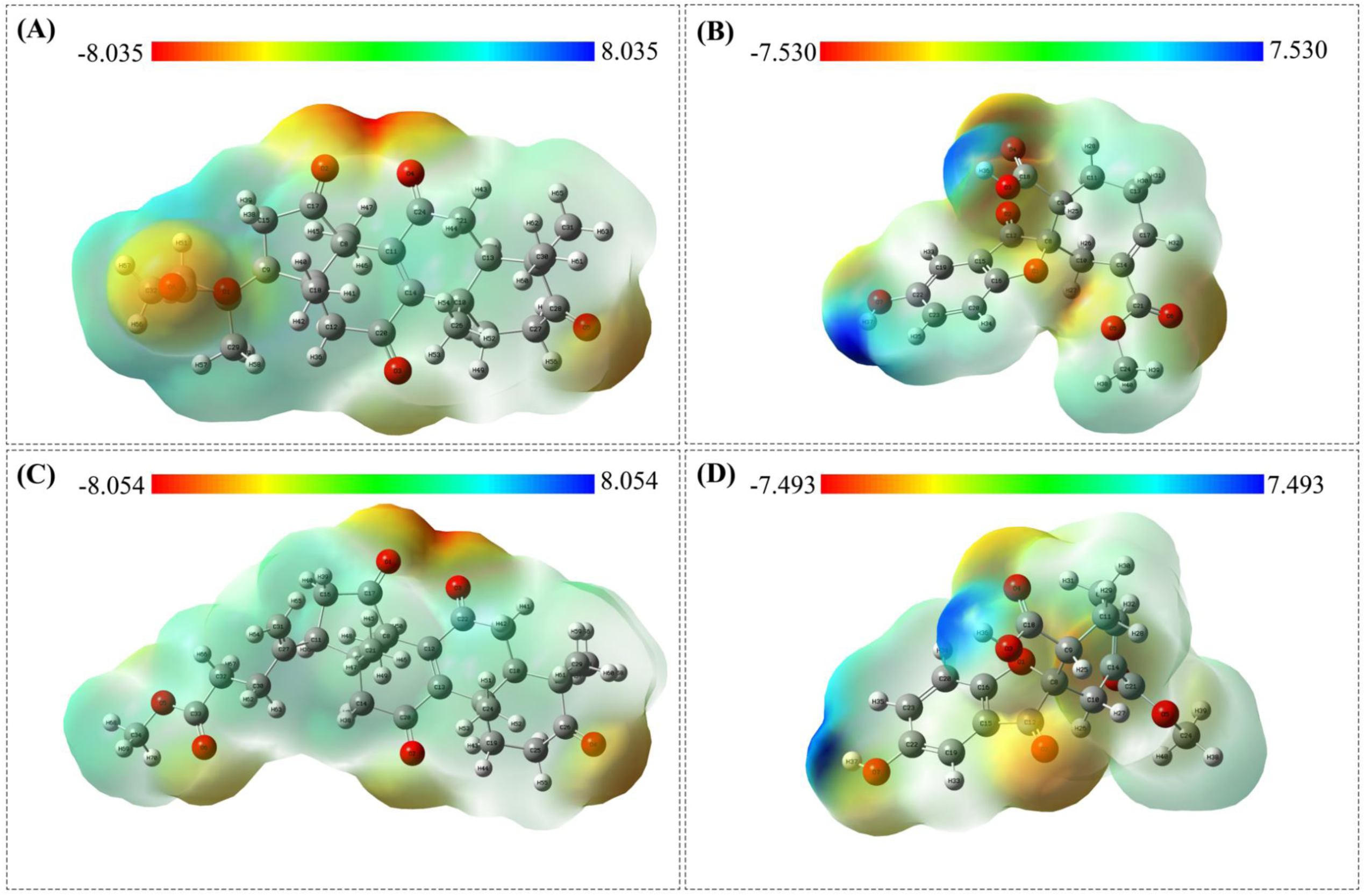
Density functional theory-derived molecular electrostatic potential surface distributions of the investigated ligands: CIDs: (A) 122184973, (B) 132962217, (C) 139584993, and (D) 132962216. The colour gradient from red to blue denotes regions of maximum negative to maximum positive electrostatic potential, respectively.

### 3.7. QSAR analysis

The QSAR prediction results showed that the selected compounds (CIDs: 122184973, 132962217, 139584993 and 132962216) have several biologically relevant activities related to antiparasitic action (**Table 5**). Among the directly related activities, CID_122184973 showed the highest predicted anti-leishmanial activity, along with good oxidoreductase inhibitory and apoptosis agonistic properties, indicating potential disruption of metabolic pathways and cell survival in the parasite. CIDs 132962217 and 132962216 were mainly characterised by their strong inhibitory effects on membrane permeability and agonistic effects on membrane integrity, suggesting potential interference with the parasites’ membrane function. Moreover, both compounds exhibited medium anti-leishmanial and anti-helminthic properties. The direct anti-leishmanial activity was comparatively low for CID_139584993. Its remarkable Myc-inhibitory, apoptosis-agonizing, and NF-κB-stimulatory properties, however, suggest an alternative mechanism that could contribute to antiparasitic activity. In general, the QSAR results indicate that the investigated compounds may exhibit antiparasitic activity through distinct and complementary mechanisms, including enzyme inhibition, induction of apoptosis, and membrane disruption.

**Table 5.** PASS-based QSAR prediction of antiparasitic-related biological activities of the selected compounds (CIDs: 122184973, 132962217, 139584993, and 132962216).

| PubChem CIDs | Pa | Pi | Activity |
| --- | --- | --- | --- |
| CID_122184973 | 0.580 | 0.016 | Antiprotozoal (Leishmania) |
|  | 0.600 | 0.037 | Oxidoreductase inhibitor |
|  | 0.672 | 0.018 | Apoptosis agonist |
|  | 0.350 | 0.030 | Antiprotozoal |
|  | 0.365 | 0.028 | DNA polymerase I inhibitor |
|  | 0.473 | 0.076 | General pump inhibitor |
| CID_132962217 | 0.409 | 0.045 | Antiprotozoal (Leishmania) |
|  | 0.330 | 0.082 | Anti-helminthic (Nematodes) |
|  | 0.763 | 0.017 | Membrane permeability inhibitor |
|  | 0.778 | 0.041 | Membrane integrity agonist |
|  | 0.541 | 0.043 | General pump inhibitor |
|  | 0.472 | 0.034 | Histidine kinase inhibitor |
|  | 0.444 | 0.025 | H<sup>+</sup>-exporting ATPase inhibitor |
| CID_139584993 | 0.794 | 0.002 | Myc inhibitor |
|  | 0.747 | 0.011 | Apoptosis agonist |
|  | 0.686 | 0.005 | Beta-glucuronidase inhibitor |
|  | 0.667 | 0.005 | Transcription factor NF-kappa B stimulant |
|  | 0.667 | 0.005 | Transcription factor stimulant |
|  | 0.637 | 0.007 | Polarisation stimulant |
|  | 0.426 | 0.039 | Antiprotozoal (Leishmania) |
| CID_132962216 | 0.763 | 0.017 | Membrane permeability inhibitor |
|  | 0.778 | 0.041 | Membrane integrity agonist |
|  | 0.761 | 0.027 | Chlordecone reductase inhibitor |
|  | 0.709 | 0.007 | Pin1 inhibitor |
|  | 0.716 | 0.032 | CYP2H substrate |
|  | 0.717 | 0.049 | Mucous-membranous protector |
|  | 0.409 | 0.045 | Antiprotozoal (Leishmania) |

## 4. Discussion

The global burden of schistosomiasis and the heavy reliance on praziquantel as the only widely used drug for treating schistosomiasis make the need for new anti-schistosomal drugs with novel targets and mechanisms of action highly critical. Current control strategies are effective against adult worms but still encounter problems due to decreased susceptibility, lack of activity against juvenile parasites and regular reinfection [1,5,47]. Thus, the discovery of novel drug candidates against key parasite proteins is a pressing challenge. In the present study, an integrated computational drug discovery approach was used to identify fungal secondary metabolites that inhibit the selected target proteins (6UY4 and 4I07) associated with *Schistosoma* species.

Natural products have been a rich source of therapeutic agents and have contributed to the discovery of many antimicrobials, antiparasitic, and anticancer drugs over the years. In particular, fungal secondary metabolites exhibit a remarkable structural diversity and biological versatility and are interesting candidates for antiparasitic drug discovery [22,23,48]. Several clinically important drugs, including penicillin G and cephalosporin C as founding β-lactam antibiotics, cyclosporin A as an immunosuppressant from *Tolypocladium inflatum*, lovastatin as a cholesterol-lowering statin from *Aspergillus terreus*, griseofulvin as an antifungal agent, and the echinocandin class of antifungals exemplify the therapeutic potential of fungal metabolites [49–52]. Recent studies have demonstrated the emerging significance of natural product-based computational screening methods in identifying lead compounds for neglected tropical diseases [48,53,54]. In accordance with these findings, the current study identified several fungal metabolites with favorable pharmacokinetic properties and strong interactions with the investigated protein targets.

CADD is becoming increasingly successful; computational methods have proven useful and reliable for identifying and optimizing therapeutic candidates. In the discovery of clinically validated drugs, integrated pipelines based on molecular docking, structure-based virtual screening, pharmacophore modeling, and molecular dynamics simulations have helped discover drugs for parasitic, bacterial, and viral diseases. For instance, the clinical efficacy of the Phase IIa clinical trial for malaria caused by *Plasmodium falciparum* (*P. falciparum*) and *P. vivax* has been obtained for DSM265 (CID_51347395), which was developed through structure-guided optimization of inhibitors against dihydroorotate dehydrogenase from the parasite [55,56]. Likewise, structure-based design has been used to develop HIV-1 protease inhibitors such as saquinavir, indinavir, and ritonavir [57,58], and docking-based virtual screening has been used to identify new candidates for the treatment of *Mycobacterium tuberculosis*, such as cefpodoxime and lymecycline [59]. Together, these studies illustrate the predictive utility and translatability of CADD and, as such, the computational pathway used in this study is a sound approach for prioritizing promising lead compounds for experimental validation.

The first step of the ADMET filtering was highly effective at reducing the library size while retaining compounds with desirable pharmacological properties, such as high gastrointestinal absorption, acceptable molecular weight, compliance with Lipinski’s rule, and the absence of major toxicity liabilities. With compounds characterized as having desirable ADMET properties, there is a higher likelihood of success in moving the compound forward to experimental validation and clinical trials [18,28–30] and thus the importance of early-stage pharmacokinetic screening is evident. The 122 compounds identified as meeting these criteria suggest that fungal metabolites are a promising source of drug-like molecules for antiparasitic drug development.

Among these molecules, CID_122184973 and CID_132962217 showed strong binding affinity towards both targets investigated. The single compound binding to several parasite proteins may increase treatment effectiveness and prevent resistance. Recent computational-directed investigations targeting Schistosoma proteins identified broad-spectrum-binding drugs as interesting lead candidates [24,54,60]. Due to their high binding affinities, the metabolites in this investigation offer medicinal potential. A thorough investigation of protein-ligand interactions revealed many hydrogen bonds, π interactions, and van der Waals contacts in both proteins’ binding sites. ASN139, GLY142, CYS141, ASN272, ASN205, and LYS241 were repeated stabilising residues in 6UY4, whereas GLN25, TRP223, HIS201, CYS31, and GLY74 were significant interacting residues in 4I07. Hydrogen bonding and hydrophobic interactions in protein-ligand complexes affect binding selectivity and affinity [24]. Multiple interactions at the active sites suggest stable and favourable binding conformations for specific fungal metabolites. Recent docking-based drug development studies have shown comparable interactions when inhibiting parasite enzymes [60].

MD simulations were used to evaluate the dynamic stability of the protein-ligand complex under near-physiological conditions. All complexes reached equilibrium quickly and remained stable throughout the 100 ns simulation, as indicated by the protein backbone RMSD analysis. The RMSD values ranged from 1.2 to 1.7 Å, indicating little structural changes due to ligand interaction. Notably, CID_132962217 had consistent RMSD values across both proteins, confirming protein-binding behaviour. In general, steady RMSD curves indicate protein-ligand compatibility and conformational equilibrium [35,61–63]. The ligand RMSD analysis was used to assess binding duration and ligand mobility in the active site. CID_132962217 showed very low RMSD values throughout the simulation, suggesting that the molecule had barely migrated from its original binding site and had good binding affinity. In contrast, CID_122184973 and CID_132962216 exhibited larger 4I07 system alterations, indicating greater ligand mobility and likely conformational changes during the simulation. The docking observations and lead compound potential of CID_132962217 are supported by its enhanced stability. Previous investigations found that compounds with low ligand RMSD had significant binding persistence and good inhibitory potential [35,61]. RMSF study assessed residue flexibility. Throughout the simulation, both proteins had modest RMSF values, indicating that ligand binding did not significantly affect their structures. Most of the highly flexible residues were in terminal and loop regions rather than in core structural domains or binding sites. Interestingly, the 4I07_132962217 complex has the lowest mean RMSF (0.96 Å), indicating higher molecular stiffness. In computational investigations of stable protein-ligand complexes [63–65], RMSF profiles are comparable and usually connected with less conformational instability. Further evidence of structural compactness and folding stability came from Rg analysis. The CID_132962217 complex has an average Rg of 17.762 Å, with minimal variation of 0.060 Å, suggesting optimal structural compactness. The Rg profile of all 6UY4 complexes remained largely constant throughout the simulation, indicating that the interaction did not trigger protein unfolding or substantial conformational changes. Long MD simulations, with or without fluctuations, show high structural integrity and protein folding, as indicated by Rg values [35,65]. The compounds’ RMSD, RMSF, and Rg studies agree that CID_132962217 is the most stable ligand.

The electronic properties that control molecular reactivity and protein interaction potential were explored using DFT analysis. Among the selected compounds, CID_139584993 exhibited the smallest HOMO-LUMO energy gap (3.552 eV) and hardness (1.776 eV), indicating relatively high chemical reactivity and electron-transferring potential. The energy gap of CID_132962216 was the highest (4.133 eV), suggesting it has high electronic stability. Conceptual DFT theory suggests that small HOMO-LUMO gaps promote electron-transfer reactions and are associated with higher biological activity and greater molecular interactions [38,66]. The relatively low hardness and high softness values of CID_139584993 and CID_122184973 thus suggest their suitability for molecular recognition and binding applications.

Furthermore, the electrostatic potential mapping revealed significant charge separation on the surfaces of the selected compounds. The electrostatic potential range of CID_139584993 was the widest, spanning -8.054 to +8.054 au, indicating a strong potential for electrostatic interactions with amino acid residues in the binding pocket. In addition, electrostatic complementarity is a key factor in enhancing protein-ligand affinity, as it allows for favorable hydrogen bonding and polar interactions [38,46]. The ESP results, therefore, support the docking and DFT results in terms of mechanism.

Based on QSAR analysis, the metabolites studied may exhibit antiparasitic activity through several synergistic biological mechanisms. CID_122184973 exhibited high activity against Leucocytozoon and anti-leishmanial activity, along with oxidoreductase-inhibitory and apoptosis-agonistic properties. CIDs: 132962217 and 132962216 exhibited significant inhibitory effects on membrane permeability and agonistic effects on membrane integrity, indicating their potential to disrupt membrane homeostasis in parasites. CID_139584993 had high probabilities of inhibiting Myc and inducing apoptosis. Multi-target activity profiles are especially beneficial in the development of anti-parasitic drugs as they decrease the risk of compensatory resistance mechanisms and can enhance therapeutic effectiveness [67,68]. These results therefore indicate that the identified compounds could act on parasites in several synergistic ways.

The results from docking, molecular interaction analysis, MD simulation, DFT calculations, ESP mapping and QSAR prediction consistently show that CID_132962217 and CID_122184973 are the most promising lead compounds. Among them, CID_132962217 showed excellent dynamic stability, good protein-binding properties, very good compactness, and biologically relevant predictions of anti-parasitic activity. Collectively, the data from these computational techniques greatly enhance confidence in the therapeutic value of these fungal metabolites.

However, there are some caveats to note. The current study is computational in nature and should be validated experimentally through enzymatic inhibition assays, parasite viability studies, and *in vivo* efficacy tests. Finally, the computational predictions are promising, but biological activity can only be inferred from *in silico* analysis. Whilst the identified compounds are promising, future studies incorporating biochemical validation, transcriptomics, and animal infection models will be crucial to assess their effectiveness against schistosomes.

## 5. Conclusion

Spiroapplanatumine F (CID_132962217) and Ganoderlactone B (CID_122184973) were shown to be potential fungal-derived dual-target inhibitors of *Schistosoma mansoni* DHODH and cathepsin. Both compounds demonstrated good docking affinities, protein-ligand interactions, favorable ADMET predictions, and stability during MD simulations. CID_132962217 had good conformational stability, while CID_122184973 had the best MM-PBSA binding profile across both targets. DFT, electrostatic potential, and QSAR bolstered their chemical reactivity and antiparasitic potential. This makes these molecules promising lead scaffolds for developing anti-schistosomal medications. These computational discoveries need enzyme inhibition, parasite viability, toxicity, pharmacokinetic, and *in vivo* research.

## Acknowledgement

The authors gratefully acknowledge the Department of Microbiology of “Primeasia University, Dhaka-1213, Bangladesh” for their assistance and support in this study.

## Author contributions

**Conceptualisation:** Tamal Paul, Swagato Dutta

**Data curation:** Md. Al-Amin, Easin Al Riad, Ananya Majumder, Aminun Naher, Fubliha Firose Auishe

**Formal analysis:** Easin Al Riad, Ananya Majumder, Aminun Naher, Fubliha Firose Auishe, Arnob Biswas, Kaniz Fatema

**Investigation:** Tamal Paul, Swagato Dutta, Sabrin Bashar

**Methodology:** Tamal Paul, Md. Al-Amin, Uma Shill, Swagato Dutta, Nowshin Tabassum

**Project Administration:** Fariha Bushra Khan, Foisal Ahmad, Khandokar Fahmida Sultana

**Resources:** Hossain Bin Afridy, Easin Al Riad, Md. Ashik Iqbal, Milon Kumar Sarkar, Arnob Biswas

**Software:** Md. Al-Amin, Aminun Naher, Ananya Majumder, Fubliha Firose Auishe, Arnob Biswas

**Supervision:** Fariha Bushra Khan, Foisal Ahmad, Ahmad Abdullah Mahdeen, Khandokar Fahmida Sultana, Sabrin Bashar

**Validation:** Tamal Paul, Md. Al-Amin, Milon Kumar Sarkar, Hossain Bin Afridy, Md. Ashik Iqbal, Sabrin Bashar, Kaniz Fatema, Arnob Biswas

**Visualisation:** Tamal Paul, Milon Kumar Sarkar, Ananya Majumder, Aminun Naher, Fubliha Firose Auishe, Kaniz Fatema, Arnob Biswas

**Writing original draft:** Tamal Paul, Md. Al-Amin, Uma Shill, Nowshin Tabassum, Swagato Dutta

**Writing-review and editing:** Tamal Paul, Ahmad Abdullah Mahdeen, Swagato Dutta, Sabrin Bashar

## Ethical statement

No ethical statement applies to this study.

## Data availability

Data are contained within the article and the supplementary information.

## Declaration of generative AI

The authors declare that ChatGPT 5.2, Grammarly, and QuillBot were utilised to assist with language editing, grammar correction, sentence restructuring, and improvement of clarity and readability. All scientific content, analyses, and interpretations presented in this manuscript are the sole responsibility of the authors.

**Supporting Information**

Supplementary File 1: ADMET analysis

Supplementary File 2: Docking results

